# CryoAPEX - an electron tomography tool for subcellular localization of membrane proteins

**DOI:** 10.1101/522482

**Authors:** Ranjan Sengupta, Michael J. Poderycki, Seema Mattoo

## Abstract

We describe a method, termed cryoAPEX, that couples chemical fixation and high pressure freezing of cells with peroxidase-tagging (APEX) to allow precise localization of membrane proteins in the context of a well-preserved subcellular membrane architecture. Further, cryoAPEX is compatible with electron tomography. As an example, we apply cryoAPEX to obtain a high-resolution three-dimensional contextual map of the human Fic (filamentation induced by cAMP) protein, HYPE/FicD. HYPE is a single pass membrane protein that localizes to the endoplasmic reticulum (ER) lumen and regulates the unfolded protein response. Alternate cellular locations for HYPE have been suggested. CryoAPEX analysis shows that, under normal/resting conditions, HYPE localizes robustly within the subdomains of the ER and is not detected in the secretory pathway or other organelles. CryoAPEX is broadly applicable for assessing both lumenal and cytosol-facing membrane proteins.

**Summary statement:** CryoAPEX couples localization of peroxidase-tagged membrane proteins at high-resolution with 3D structural analysis, within an optimally preserved cellular context.

## Introduction

Localization of membrane proteins via electron microscopy (EM) at high resolution is dependent on robust detection technology and on sample preparation methods that confer superior ultrastructural preservation of membranes. Unfortunately, current methods of localization of membrane-bound proteins at EM resolutions are less than optimal. Immunoelectron microscopy (IEM) to detect either an endogenous or epitope-tagged overexpressed protein using antigen-specific antibodies requires a permeabilization step that also causes degradation of cellular membranes and distortion of membrane-bound compartments *(De Mey, 1981; Schnell et al., 2012)*. An alternative is to fuse enzymatic tags directly to the protein of interest in a transfection experiment, so as to avoid the necessity for introducing an antibody. A number of these enzymatic tags have been described, e.g., metallothioneine (METTEM), resorufin arsenical hairpin (ReAsH), miniSOG, horse radish peroxidase (HRP) and more recently, engineered ascorbate peroxidases (APEX and APEX2) *(Hoffmann et al., 2010;* Lam *et al*., 2015; *Martell et al., 2012*; *Mercogliano et al., 2007; Porstmann et al., 1985; Shu et al., 2011)*. While each of these technologies faces its own set of limitations when employed in conjunction with EM, the APEX2 tag, which has been used with success on mitochondrial and ER proteins, is arguably the most promising option *(Martell et al., 2017)*. APEX2 is a monomeric 28kDa soyabean ascorbate peroxidase that withstands strong EM fixation *(Lam et al., 2015)*. Additionally, it is sensitive, generally straightforward in its application and unlike horseradish peroxidase is active in both the cytosolic and lumenal compartments *(Hopkins et al., 2000; Lam et al., 2015; Martell et al., 2017)*. Nevertheless, the morphological damage to cellular membranes and membrane-bound organelles that occurs during conventional aldehyde fixation/alcohol dehydration protocols, even without the permeabilization required for antibodies, continues to be an impediment to obtaining optimal preservation of the subcellular architecture.

In contrast, cryofixation or high pressure freezing (HPF) is a method for obtaining vitreous preparations of live cells and tissues up to 200μm in thickness with minimal ice crystal formation, thus immobilizing macromolecular assemblies in their near-native state *(Chan et al., 1993; Mcdonald 1999; Zechmann et al., 2007; Studer et al., 2008)*. This method has become a mainstay for preparing samples for electron tomography, which employs thicker sections *(McDonald et al., 2006)*. Further, HPF has been adapted in combination with freeze substitution (FS) methods, which entail organic substitution of water with acetone at low temperature, to generate plastic embedded samples for conventional EM. However, HPF/FS has not been extensively used with protein localization methods that require chemical fixation of cells (*Tsang et al., 2018*).

Here, we opted to develop an EM tomography compatible detection method to visualize the human Fic (filamentation induced by cAMP) protein, HYPE/FicD. Fic proteins are a recently characterized class of enzymes that predominantly utilize ATP to attach AMP (adenosine monophosphate) to their protein targets *(Casey & Orth, 2018; Truttmann et al., 2017; Worby et al., 2009)*. This post-translational modification is called adenylylation or AMPylation. The first Fic proteins, VopS and IbpA, were described in the pathogenic bacteria *Vibrio parahemolyticus* and *Histophilus somni*, respectively, where they serve as secreted bacterial effectors that induce toxicity in host cells by adenylylating/AMPylating and inactivating small GTPases *(Mattoo et al., 2011; Worby et al., 2009; Xiao et al., 2010; Yarbrough et al., 2009; Zekarias et al., 2010)*. Fic proteins have also been implicated in bacterial cell division and persister cell formation, protein translation, cellular trafficking, and neurodegeneration *(Garcia-Pino et al., 2014; Harms et al., 2015; Mukherjee et al., 2011; Truttmann et al., 2018)*.

HYPE (huntingtin yeast interacting protein E) or FicD is the sole Fic protein encoded by the human genome (*Faber et al., 1998; Sanyal et al., 2015)*. In humans, HYPE is expressed ubiquitously, albeit at very low levels. It is a single pass type II membrane protein that localizes to the lumenal surface of the endoplasmic reticulum (ER) *(Sanyal et al., 2015; Worby et al., 2009)*. HYPE plays a critical role in regulating ER homeostasis by reversibly AMPylating the Hsp70 chaperone, BiP/GRP78 *(Ham et al., 2014; Sanyal et al., 2015; Preissler et al., 2015; Preissler et al., 2017a; Preissler et al., 2017b)*. Biochemical and proteomic screens have identified additional AMPylation targets of HYPE and its orthologs, which include cytosolic chaperones, cytoskeletal proteins, transcriptional and translational regulators, and histones *(Broncel et al., 2016; Truttmann et al., 2016; Truttmann et al., 2017)*. These data suggest that a fraction of HYPE could reside outside the ER, e.g. in the nucleus or cytoplasm. Indeed, a small fraction of the HYPE homolog in *C. elegans*, Fic-1, has been shown to localize to the cytosol and AMPylate cytosolic Hsp70 and Hsp40 proteins *(Truttmann et al., 2017)*.

The low levels of HYPE expression in human cells combined with the resolution limitations of conventional immunofluorescence microscopy make obtaining definitive localization data difficult. We, therefore, opted to circumvent these limitations by using an electron microscopy approach. To visualize HYPE in cells, our challenge was to develop a technique that would preserve membrane ultrastructure and be compatible with transmission EM tomography methods to identify specific distribution of HYPE in three-dimensional (3D) space. We chose to visualize HYPE by genetically tagging it with APEX2. The APEX2 tag catalyzes a peroxide based reaction that converts DAB (diaminobenzadine) into a low-diffusing precipitate that deposits at the site of the target protein *(Lam et al., 2015)*. In our analysis of HYPE, we also included another protein, ERM (endoplasmic reticulum membrane), to serve as a dual control for ER localization as well as for ER morphology. ERM consists of the N-terminal 1-27 amino acids of the P450 oxidase 2C1 (*Lam et al., 2015)*. An ERM-APEX2 fusion localizes to the ER membrane such that the APEX tag faces the cytosol. Additionally, ERM is known to induce a reorganization of the smooth ER into distinctive ordered membrane structures called OSERs (Organized Smooth ER) (*Snapp et al., 2003; Lam et al., 2015; Sandig et al., 1999)*. Thus, ERM-APEX2 serves as an excellent metric for assessing both ER membrane specific staining and ultrastructural membrane preservation.

Next, since degradation of the cellular ultrastructure in traditional aldehyde fixation/alcohol dehydration methods appears to be largely associated with the alcohol dehydration post-processing steps and not with aldehyde fixation per se, we adopted and optimized a combination method proposed by Sosinsky *et al*., which relies on chemical fixation prior to cryofixation and optimally preserves membrane structure *(Sosinsky et al., 2008)*. We, therefore, applied this combination approach towards the detection of APEX2 tagged proteins. Using this methodology, we observed minimal lipid extraction or distortion of membrane structures, and were able to clearly detect ER membrane-bound HYPE. During submission of this manuscript, a similar conceptual approach of combining HPF and APEX tagging, albeit procedurally different and at a lower resolution, was reported for correlative light-electron microscopy (CLEM) on whole tissues *(Tsang et al., 2018)*. We show that the addition of tannic acid, uranyl acetate and counter staining with lead further increased the overall contrast, and made the membrane ultrastructure in the vicinity of the localized HYPE-APEX2 density easily recognizable. Such optimizations proved key to making this technology amenable to transmission EM tomography of ER membranes.

Our hybrid method - hereon referred to as cryoAPEX - performed remarkably well for protein localization at the subcellular level. Comparison of ERM-APEX2 in cryoAPEX treated cells versus live cryofixed (HPF alone) cells showed well-preserved OSER morphology with ER specific staining only on the cytosolic face of the ER membrane. Similarly, we observed that HYPE localizes as periodic foci along the lumenal face of the ER membrane and, although overexpressed in the context of a transient transfection study, is retained only within the ER and the contiguous nuclear envelope. HYPE remained ER associated even when the ER came in close proximity to the plasma membrane, mitochondria or the Golgi apparatus. Thus, it would appear unlikely that HYPE traffics via conventional means through the secretory system or is associated with other organelles under normal circumstances. Further, data collected on cryoAPEX treated cells could be used to reconstruct a 3D representation of HYPE within the ER lumen. This ability to simultaneously assess the detection of a membrane protein in multiple cellular compartments throughout the subcellular volume of a single cell at low-nanoscale resolution is a significant advance. More broadly, we present a straightforward methodology for probing the subcellular distribution of membrane-bound proteins, with either lumen-facing or cytosol-facing topologies, that is amenable to high-resolution 3D tomographic reconstruction.

## Results

### A combination of glutaraldehyde fixation, cryofixation and extended osmication during freeze substitution shows optimal preservation of membrane ultrastructure in HEK-293T cells

In cellular imaging, the ability to obtain 3D spatial localization of proteins in the context of well-preserved cellular structures at high resolution is coveted *(O’Toole et al., 2018)*. For antibody-conjugate based detection methods to be effective, ultrastructure-damaging permeabilization and/or technically challenging ultracryosectioning are required. At EM resolutions, the deleterious effects of such treatments, particularly upon membrane-bound compartments, become obvious. Fusion of a protein of interest to the monomeric enhanced peroxidase APEX2 avoids the need for an antibody. However, the published protocols still use chemical fixation and alcohol dehydration prior to embedding *(Martell et al., 2017)*. We hypothesized that the alcohol dehydration step in published APEX2 protocols is limiting for ultrastructural preservation, and consequently for signal intensity and resolution. Therefore, referencing work by Sosinsky *et al (Sosinsky et al., 2008),* we utilized cryofixation/freeze substitution (HPF/FS) of live cells as a gold standard for preservation, and combined it with chemical fixation methods to optimize ultrastructural preservation in untransfected human HEK-293T cells.

Preservation was assessed based on several criteria including membrane integrity, smoothness of intracellular membranes, densely packed cytoplasm, and maintenance of organellar structures such as mitochondria with clearly visible cristae. Smoothness of intracellular membranes is often a primary indication of good preservation, and preservation of the nuclear envelope has been used classically as a hallmark (*Sosinky et al., 2008; Tsang et al., 2018*). As shown in Figure 1, panels A-C, traditionally used glutaraldehyde fixation/alcohol dehydration methods showed poor preservation of the nuclear membrane with intermittent ruffling and separation of the double membrane. The cytoplasm of these cells also appeared less granulated and less densely packed (Supplemental Figure S1, panels A, B and G). In contrast, the HPF/FS processed samples showed excellent preservation of the nuclear envelope, which was smooth and devoid of distortion (Figure 1, panels G-I). Additionally, the cytoplasm was densely packed and the mitochondrial membrane remained intact, with clearly visible cristae (Supplemental Figure S1, panels E, F and I). Next, we fixed our cell samples with glutaraldehyde prior to HPF/FS (Figure 1, panels D-F and Supplemental Figure S1, panels C, D and H). Morphological damage beyond that seen with HPF/FS alone was minimal, and this combination approach conferred substantially better ultrastructural preservation relative to traditional fixation techniques. This excellent membrane preservation via the combination (Glut+HPF+FS) method was further substantiated when we extended it to another cell line, BHK (Baby hamster kidney) cells that are routinely used in electron microscopy studies (*Hawes et al., 2007*). Indeed, a comparison of ultrastructural preservation using HPF/FS (Figure 1, panels J-L) versus our combination method (Figure 1, panels M-O) showed similar results, as determined by a well-preserved nuclear membrane (red arrows), and comparable preservation of mitochondrial (yellow arrows) structure and ER (blue arrows) membranes.

**Figure 1:**
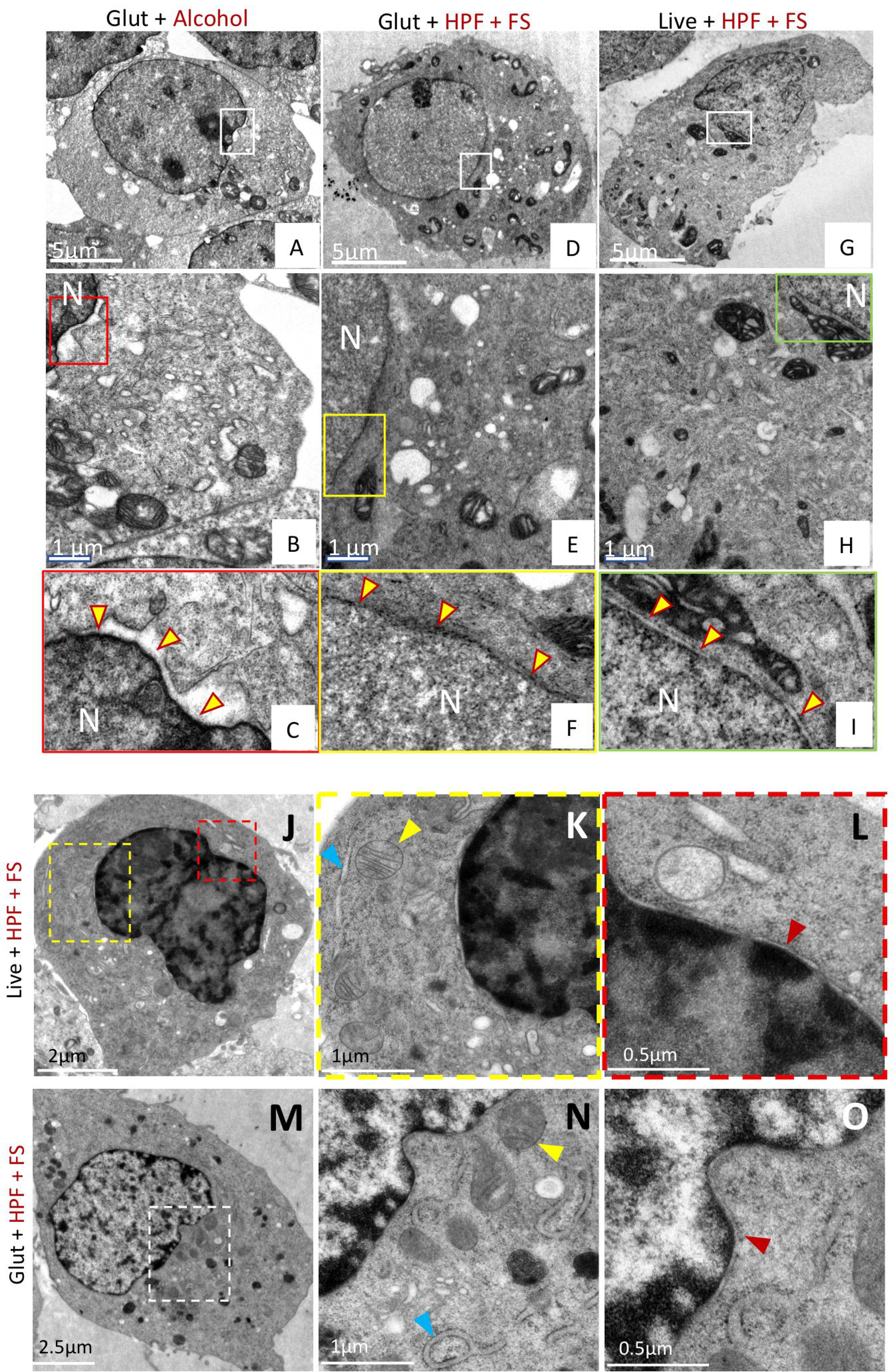
A combination of chemical fixation and cryofixation exhibits superior ultrastructure preservation and membrane staining over traditional methods in HEK-293T cells. Cells prepared by conventional glutaraldehyde fixation and dehydration methods (A-C); or glutaraldehyde followed by cryofixation (D-F); or cryofixation alone (G-I) were examined by thin-section EM. Panels B, E and H represent magnified views from boxed regions of panels A, D and G, respectively. Panels C, F, and I represent magnified views of the boxed regions from panels B, E, and H, respectively, and highlight the preservation of the nuclear membrane in each case (yellow arrows). The nuclear membrane appears smooth and uncompromised in samples prepared by glutaraldehyde/cryofixation (Glut+HPF+FS) or cryofixation (HPF+FS) alone, but is ruffled and irregular in samples prepared by conventional means (Glut+Alcohol). Glut = glutaraldehyde. HPF = high pressure freezing. FS = freeze substitution. N = nucleus.

### ERM-APEX2 induced OSER structures show that membrane preservation is improved with the cryoAPEX method

Next, we sought to test the applicability of the above combination method for ultrastructural preservation with the use of the APEX2 tag to follow a protein of interest. Hereon, we refer to this hybrid glutaraldehyde+HPF/FS+APEX2 method as cryoAPEX (Figure 2-1, flowchart).

**Figure 2:**
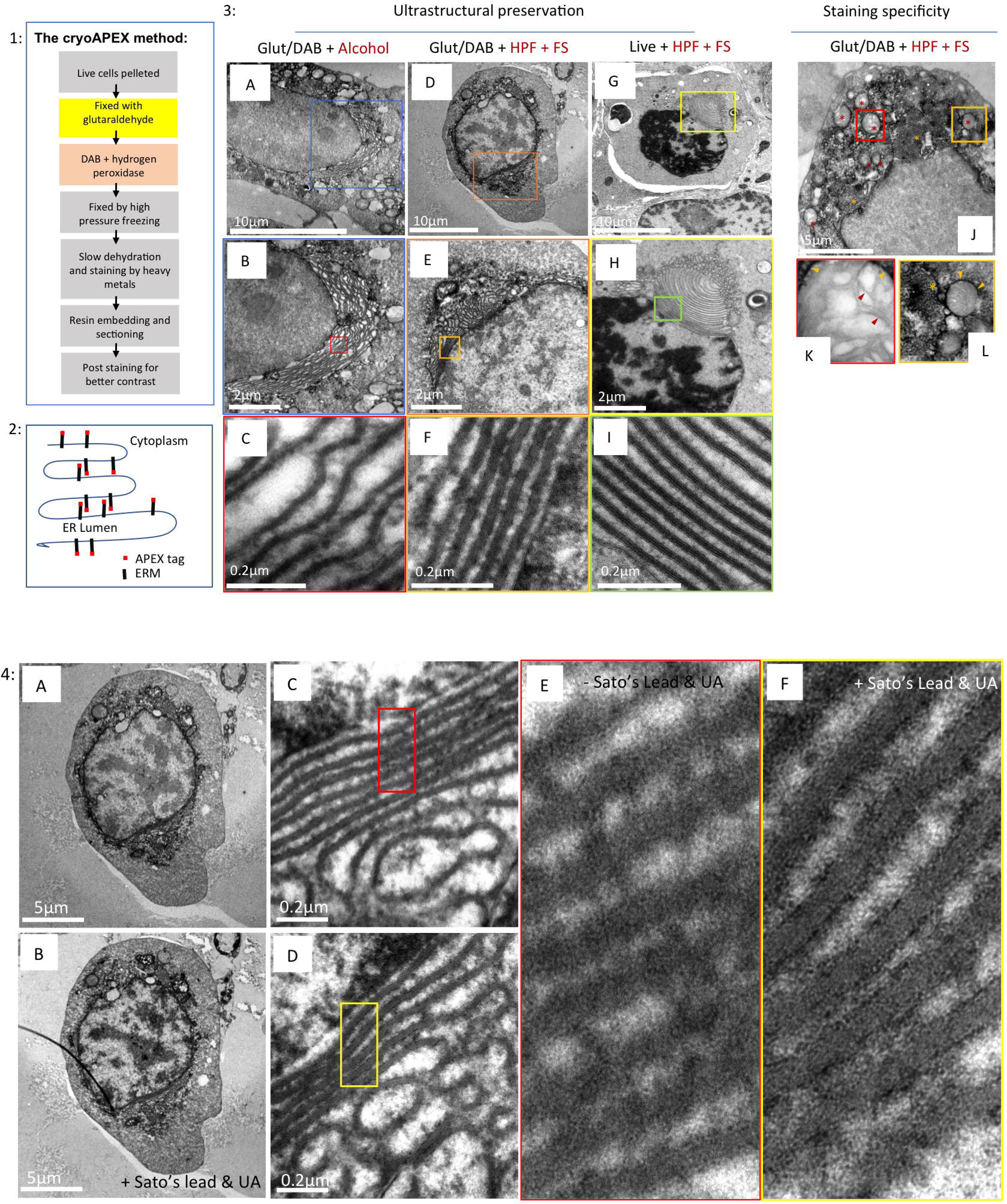
OSER as a system for evaluating membrane preservation and staining specificity. **1)** Flowchart describing cryoAPEX. **2)** Schematic of APEX-tagged ERM expressed on the cytosolic face of the ER membrane. **3)** The reorganized ER morphology in chemically fixed, DAB reacted ERM-APEX2 expressing cells that were processed via traditional chemical fixation/alcohol dehydration (A-C) or by cryoAPEX (D-F) was compared to ERM-APEX2 expressing cells that were cryofixed live and without the DAB reaction (G-I). The live cryofixed cells (G-I) represent the best attainable ultrastructural preservation and serve here as the metric for evaluating membrane preservation obtained via the two APEX based detection protocols (A-F). The high specificity of staining obtained by cryoAPEX is exemplified in images J-L. Here, thin section images of cells expressing ERM-APEX2 processed by cryoAPEX show preferential staining of the reorganized ER (J, orange asterisk) and the outer mitochondrial membrane (red asterisk in J and orange arrow heads in K-L, respectively) but not of the mitochondrial cristae (K, red arrow heads). **4)** Post staining with heavy metals improves definition of preferentially stained membranes. Post-staining of thin sections with heavy metals using uranyl acetate and Sato’s lead solution following cryoAPEX provides additional contrast, thereby improving resolution. Shown are thin sections of the same cell imaged in Figure 2-3D before (A, C, and E) and after (B, D, and F) post-staining. Comparison of panels E and F clearly shows improved definition and the resolution of membranes at high magnifications in post-stained samples (F). UA = uranyl acetate.

To serve as a proof of concept, we evaluated cryoAPEX by assessing the membrane localization of the ERM-APEX2 chimeric protein (Figure 2-2). ERM (Endoplasmic reticulum membrane) consists of amino acid residues 1-27 of the P450 oxidase 2C1 (*Lam et al., 2015*; *Sandig et al., 1999)*. Its expression results in ER localization of the peptide as a membrane protein, such that the peroxidase tag faces the cytoplasm *(Lam et al., 2015;* Figure 2-2). This topology in the ER membrane serves as an additional control for HYPE, which displays the opposite orientation in the membrane. ERM expression is known to cause reorganization of the smooth ER and increased membrane biogenesis *(Lam et al., 2015; Sandig et al., 1999)*. Cells expressing ERM form distinctive ordered labile membrane structures called OSERs (Organized Smooth ER) that can be easily visualized via electron microscopy without the need for specialized detection methods *(Snapp et al., 2003)*. Thus, it is possible to conveniently assess the degree of membrane preservation in conjunction with various sample preparation methods, providing an excellent metric for assessing both staining specificity and ultrastructural preservation in one system.

As a morphological control, thin sections of ERM-APEX2 transfected cells were processed using HPF/FS alone and then stained using osmium tetroxide, tannic acid and uranyl acetate but without chemical fixation or the DAB reaction. Under these conditions, the typical membrane whorl pattern of reorganized ER adjacent to the nucleus was clearly visible (Fig 2-3, panels G-I, yellow and green boxes). OSER membranes within these were arranged in evenly spaced parallel stacks without disruption or ruffling (Fig 2-3, panel I). In contrast, traditional methods showed a clear disruption of the lamellar stacking of these OSER structures (Figs. 2-3, panels A-C and supplemental Figure S2A-D), with artefactual ruffling of the membrane stacks and presumed loss of cytoplasmic material between the stacks (compare Fig 2-3C and supplemental Fig S2D with 2-3I). However, the cryoAPEX method (Figs. 2-3, panels D-F and supplemental Figure S2, panels E-H) resulted in well preserved ER derived lamellar structures comparable to the lamellar stacking in ER derived structures observed in live HPF/FS controls (compare Fig 2-3F and supplemental Fig S2H with 2-3I). Peroxidase-reacted samples were also processed in parallel without uranyl acetate, since uranyl acetate stains nucleic acids and therefore the nucleus and to some extent, the mitochondria (Figs, 2-3, panels J-L, red asterisks). Tannic acid was also eliminated from these samples to minimize background staining (Figs. 2-3, panels J-L) (Huxley et al., 1961.; Kalina and Pease, 1977; Persi and Burnham, 1981; Schrijvers et al., 1989). Staining was not observed within the nucleus (Figs. 2-3J) or in mitochondrial cristae (red arrowheads within red box) (Figs. 2-3K) but was seen in the ER derived structures (Figs. 2-3K and 2-3M, yellow asterisks and yellow arrows within yellow box). As additional controls for the specificity of the ERM-APEX2 staining for the ER, we assessed localization of three organellar markers – namely, mito-V5-APEX2 that targets to the mitochondrial matrix (Addgene plasmid #72480; *Lam et al., 2015*); CAAX-APEX2 that targets to the plasma membrane (*Lam et al., 2015*), and ManII-APEX2 that targets to the golgi lumen (See Methods). Supplemental Figures S3A, S3B, and S3C show APEX2 associated staining only at the mitochondria, golgi, and plasma membrane, respectively, confirming that APEX2 associated signals obtained in our assays are specific to the tagged protein.

To enhance the contrast of membrane straining over the background of overall osmicated DAB density, the same sections shown in Figure 2-3, panels D-F were imaged before and after post-staining with Sato’s lead solution and uranyl acetate (Fig 2-4). This resulted in an improved ability to distinguish specific staining of adjacent membranes at higher magnifications (Fig 2-4, compare panels A, C and E with B, D, and F, respectively).

### HYPE localizes to the ER membrane as distinct lumen-facing foci

Having established the methodology to localize an APEX2-tagged ER membrane protein in an optimally preserved cell, we next applied cryoAPEX to the sole human Fic protein, HYPE. Previous immunofluorescence, cell fractionation, and protease protection assays have placed HYPE as predominantly facing the ER lumen *(Sanyal et al., 2015)*. A smaller fraction of HYPE has also been detected in the cytosol, as well as in the perinuclear space *(Truttmann et al., 2015; Truttmann et al., 2017*). We, therefore, transiently transfected HEK-293T cells with HYPE-APEX2 constructs and processed them by cryoAPEX as standardized for ERM-APEX2 above. As before, ultrastructure was well preserved and there were minimal signs of lipid extraction or membrane ruffling (Figure 3). Osmicated DAB density was observed within the lumen of peripheral ER tubules (Figs. 3A and 3B, yellow and red boxes, respectively) but not in identically treated untransfected controls (Figs. 3D and 3E, yellow and red boxes). Remarkably, at higher magnification this density resolved into distinct foci along the length of the tubules, on the lumenal face of the ER membrane (Fig 3C, highlighted by white arrowheads). These HYPE-specific foci averaged a distance of 61.45nm apart (Figure 3D). Further, this density was not an artifact of the rough ER, where such density corresponding to the presence of ribosomes is seen only on the cytosolic face of the ER membrane (Fig 3G). Interestingly, we rarely observed ribosomal density on the outer face of HYPE-APEX stained ER membranes. To emphasize the merits of cryoAPEX over traditional methods, we also assessed HYPE-APEX2 transfected cells using the traditional alcohol dehydration method (Fig S4). Analysis of thin sections using this method revealed a similar staining pattern within the tubules of the peripheral ER and nuclear envelope (Fig S4B, white arrows within yellow box). However, as expected, there was poor preservation of the ER membrane and intermittent regions of membrane discontinuity (Fig S4B, red arrowheads). Additionally, the presence of other membrane-bound structures was mostly indiscernible (compare Fig S4A with Fig 3A), thus making it impossible to get contextual information about the other organelles in the immediate vicinity or in contact with the ER membrane containing HYPE. Clearly, cryoAPEX offers high resolution localization for HYPE in the ER lumen in the context of other organelles at the subcellular level.

**Figure 3:**
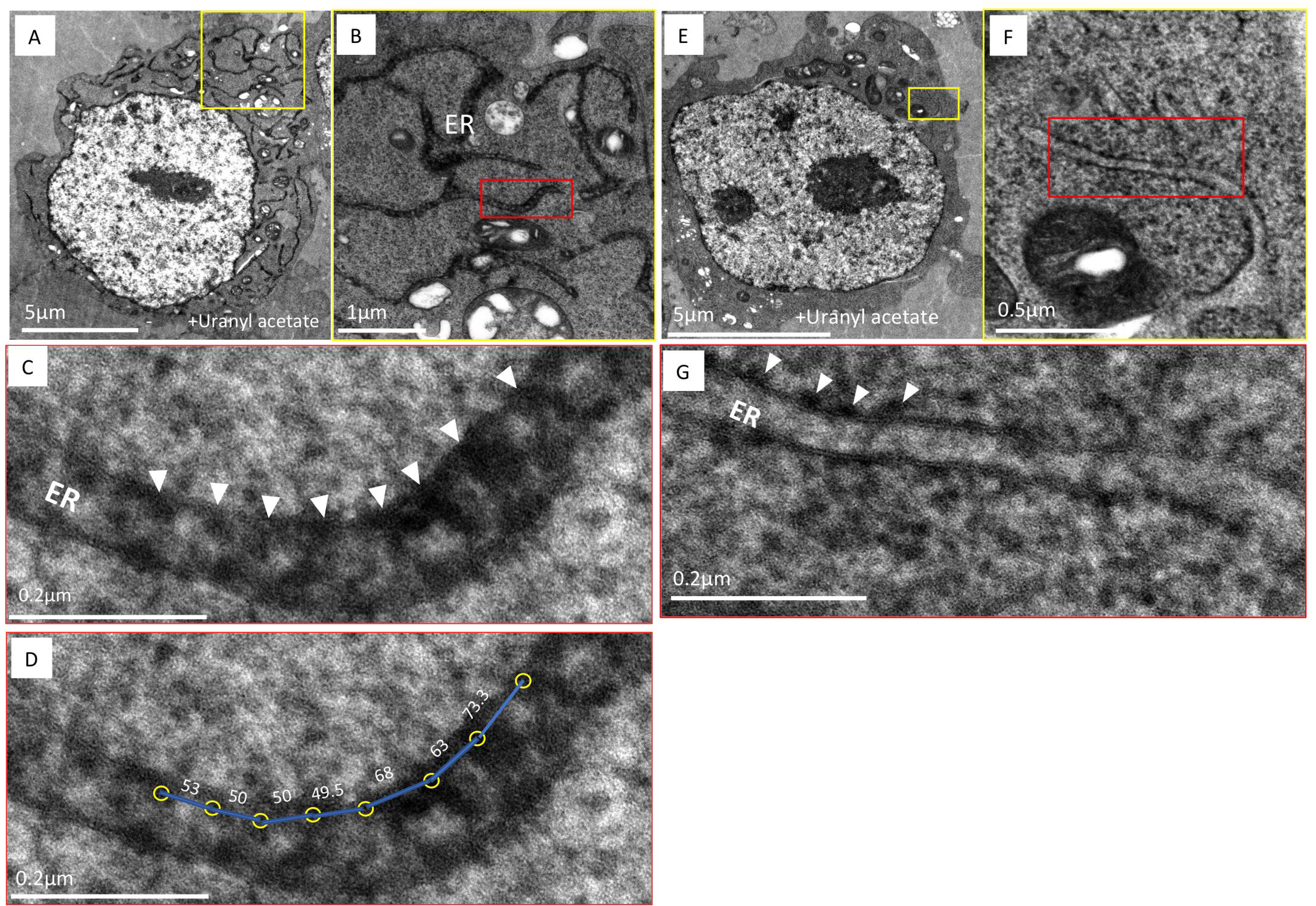
HYPE localizes to the lumenal face of the ER membrane as periodic foci. **(A)** An image of a thin section of HEK-293T cells expressing HYPE-APEX2 and processed by cryoAPEX reveal staining of the ER tubules in a well preserved (dense) cytoplasmic background. **(B and C)** Higher magnification images of a small section of the peripheral ER (demarcated by yellow box in A) exhibits periodic foci of APEX2 generated density (B, red box and C, white arrow heads showing periodicity between the HYPE foci). **(D)** Center-to-center density measurements showing the distance (in nm) between the HYPE-specific foci. **(E)** Untransfected control cells processed in an identical manner show the lack of APEX2 generated density within the ER lumen. **(F and G)** A higher magnification image of E (demarcated by yellow box) clearly shows the lack of density on the lumenal face (F, and magnified image of the ER section within red box in G). Additionally, density corresponding to ribosomes on the cytoplasmic face of the ER membrane is evident (G, white arrowheads). Thus, HYPE is detected only along the lumenal face of the ER membrane and never on the cytosolic face.

### Assessing localization of HYPE in cellular compartments other than the ER

In addition to the ER, HYPE has been suggested to interact with protein targets in the cytosol and nucleus, and possibly in the secretory pathway (*Broncel et al., 2016; Rahman et al., 2012; Truttmann et al., 2016; Truttmann et al., 2017)*. We, therefore, screened for the presence of HYPE in subcellular compartments other than the ER. HYPE was present within the lumenal walls of the NE (nuclear envelope), which is expected since the nuclear membrane is contiguous with the ER (Fig 4-1, A-C, yellow and red boxes). This is in agreement with immunofluorescence data, which shows HYPE specific staining in the perinuclear membranes *(Sanyal et al., 2015; Truttmann et al., 2015)*. The characteristic periodic foci of osmicated DAB density were seen as well (Fig 4-1, C, red arrows). As a control, untransfected cells were also processed using cryoAPEX; no lumenal staining was evident, indicating the specificity of HYPE’s NE localization (Fig 4-1, D-F, red box and white arrows in F).

**Figure 4:**
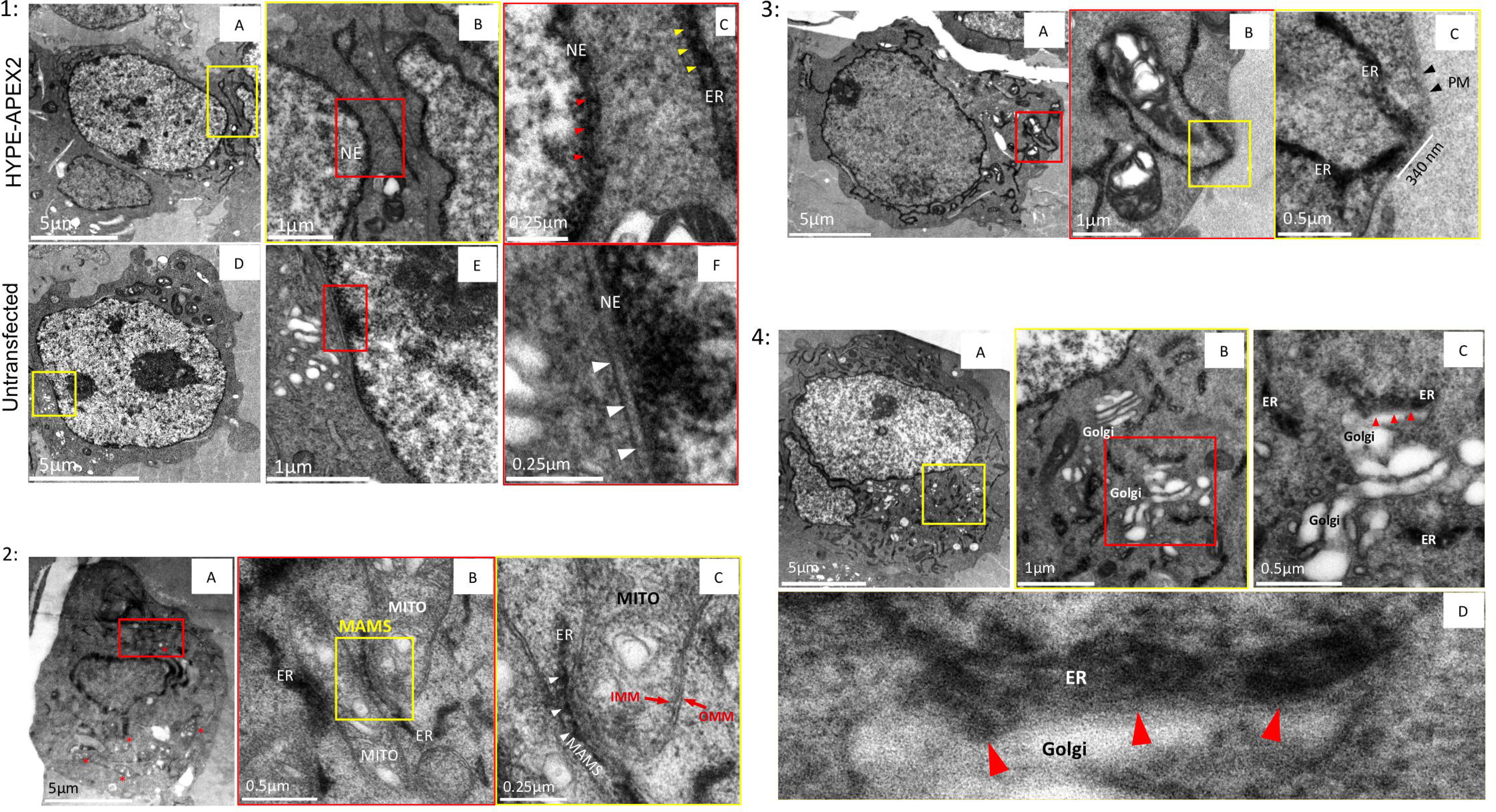
Assessing localization of HYPE in subcellular compartments other than the ER. **1) Localization of HYPE to the nuclear envelope (NE):** Images of thin sections from cells transfected with HYPE-APEX2 and processed by cryoAPEX show HYPE specific density within the perinuclear space of the nuclear envelope (A-C). At higher magnification, this staining shows a pattern similar to that seen along the lumenal face of the ER membrane (C; compare red and yellow arrowheads within the NE and the ER, respectively). Untransfected cells processed in this manner exhibit no membrane associated staining within the perinuclear space of the nuclear envelope (D-F and white arrowheads in F). **2) HYPE does not localize to the mitochondria:** Cells transfected with HYPE-APEX2 were processed by cryoAPEX in the presence of osmium tetroxide but without addition of uranyl acetate or tannic acid. Thin sections of these cells showed a distinct lack of mitochondrial membrane staining, making it difficult to visualize mitochondria at low magnifications (red asterisks in A). Magnified images of well-preserved ER-mitochondrial junctions (MAMS; demarcated by red box in A) clearly show ER tubules in close contact with two adjacent mitochondria (B). A further magnified image of the MAMS shows the HYPE-APEX2 staining of the ER but no apparent staining within the inner or outer mitochondrial membranes (B, yellow box; and C, magnified image of the area within this yellow box). The periodic distribution of HYPE is retaied even at the MAMS (C, white arrow heads). ER = endoplasmic reticulum. MITO = mitochondrion. IMM = inner mitochondrial membrane. OMM = outer mitochondrial membrane. MAMS = mitochondria associated membranes. **3) HYPE does not localize to the Plasma Membrane (PM):** Images of a cell expressing HYPE-APEX2 reveal absence of plasma membrane staining (A). Images of ER-PM junctions at high magnifications show well preserved junctions where the ER makes extended contacts with the plasma membrane (red box in A; yellow box in B; white bar in C). Staining was contained within the cortical ER tubules with no apparent staining of the plasma membrane (C, black arrowheads) **4) HYPE does not enter the secretory pathway:** To assess whether HYPE enters the secretory pathway, the Golgi apparatus was imaged (A-D). Images from thin sections of HYPE-APEX2 transfected cells show an area where ER tubules are interspersed within Golgi stacks (A; region within yellow box). Higher magnification of this region shows the typical stacked morphology of the Golgi apparatus devoid of any osmicated DAB density within its lumen (B and C). Further magnification of a well-preserved ER-Golgi junction shows a region of extensive contact between the two organelles where the DAB density was restricted to the ER lumen and showed no apparent staining of the Golgi apparatus (B, area within red box. Red arrows in C and D indicate the ER-Golgi junction).

We next investigated the presence of HYPE in the mitochondria. Samples were prepared without uranyl acetate to avoid staining mitochondrial nucleic acids. Osmicated DAB density was observed at multiple junctions between the ER and mitochondria in membranes called MAMS (mitochondria associated membranes), which are contiguous with the ER, and displayed the typical periodic HYPE foci (Fig 4-2, A-C, white arrowheads in C). Upon close examination, it was clear that although the two organelles were in close proximity, the inner and outer mitochondrial membranes were devoid of staining (Fig 4-2, C). Thus, HYPE staining remained specific for the contiguous ER and did not show any evidence for the existence of HYPE within the mitochondrion.

Similarly, we also searched for evidence of HYPE staining at the plasma membrane (PM), as the cortical ER is known to make multiple PM contacts. Even at these contact points, osmicated density was restricted to the ER lumen and was clearly absent from the adjoining PM (Fig 4-3, A-C).

Finally, we assessed the Golgi apparatus for evidence as to whether or not HYPE enters the secretory pathway. Uranyl acetate and tannic acid, in addition to osmium tetroxide, were used to enhance the contrast for the DAB density at Golgi-ER junctions (Fig 4-4, red arrows). The typical stacked morphology of the Golgi was clearly visible, interspersed with ER tubules (Figs 4-4, A-C, yellow and red boxes). Even at the Golgi-ER junction, the HYPE specific periodic DAB density was clearly restricted to the ER lumen and was completely absent from the Golgi (Fig 4-4, D).

Altogether, at the expression levels assessed here, HYPE is localized to the ER lumen and contiguous nuclear envelope, and does not appear to localize or transport to adjacent organelles or the plasma membrane even upon close contact with the ER.

### CryoAPEX allows tracking of the subcellular distribution of HYPE and ERM over a large cellular volume

The greatest advantage of using cryofixation for sample preparation for EM is its capability to preserve membrane ultrastructure consistently throughout the cell volume. To track the distribution of HYPE over a large subcellular volume, we employed serial section EM (Figure 5). Multiple such ribbons containing between 10-20 serial sections, each 90nm thick, were collected and screened. Representative images of a subset of 8 serial sections selected from a larger set of images of a demarcated region from a single cell are depicted (Figure 5A, yellow box and panels numbered 1 through 8). The images capture a region of a peripheral ER network showing osmicated DAB density within the ER and in close proximity to the mitochondria and Golgi apparatus. The excellent preservation of the ultrastructure of these neighboring organelles and the additional uranyl acetate staining provide contextual information within which HYPE resides. A similar analysis of ERM-APEX2 is shown in Fig S5. Thus, such preservation and staining using cryoAPEX allows us to subsequently use EM tomography to determine the localization of membrane proteins like HYPE and ERM in 3D.

**Figure 5:**
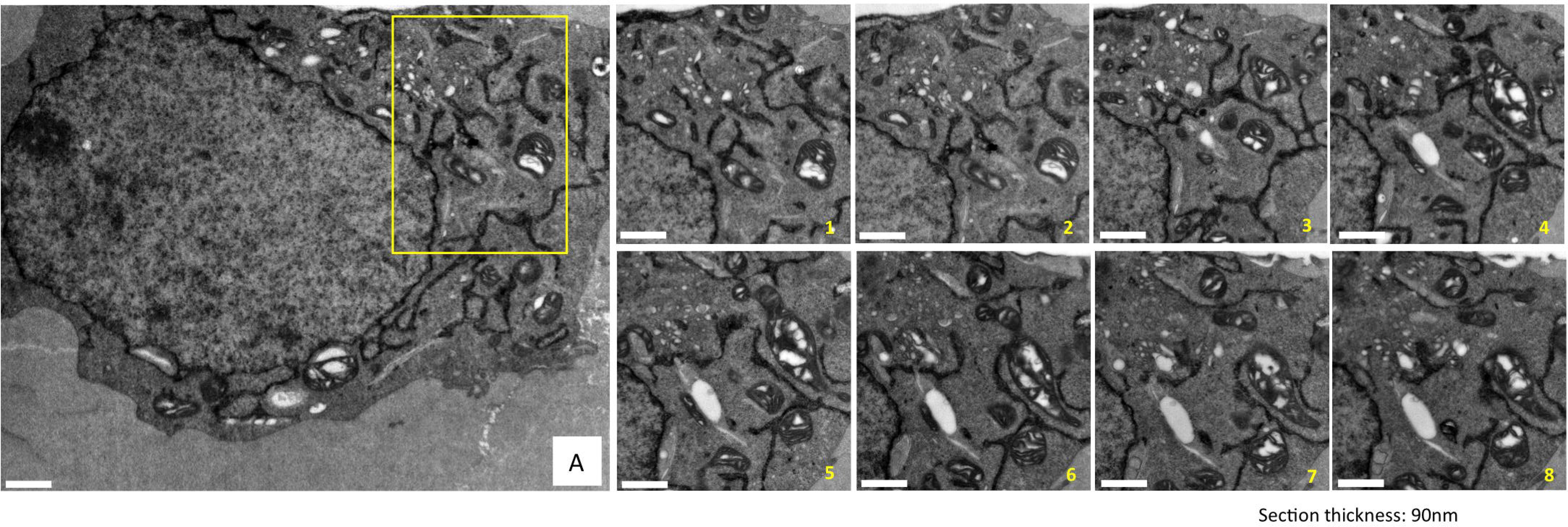
Superior ultrastructural preservation enables the tracking of HYPE’s subcellular localization via serial sectioning. To demonstrate the consistency of the membrane ultrastructure preservation obtained by cryoAPEX, cells expressing HYPE-APEX2 were serially sectioned and a specific area (panel A, yellow box) was imaged. Multiple such ribbons containing between 10 and 20 serial sections of 90nm thickness were collected, screened and imaged. Representative images of 8 serial sections showing ER localization of HYPE are presented (images serially numbered 1 though 8). Sections exhibit a dense well-preserved cytoplasm with undisrupted membrane ultrastructure of organelles like mitochondria and Golgi complex in close proximity to the ER tubules containing HYPE-APEX2 density. Thus, we can follow HYPE localization through the volume of the cell without loss of contextual information.

### EM tomographic reconstruction of the ER from a cell expressing HYPE-APEX2

To visualize HYPE, we collected a tilt series from a single 250nm section of HEK-293T cells expressing HYPE-APEX2 (Fig 6A-1). The 3D reconstruction of HYPE within the ER was carried out for the region of the cell demarcated by the red box (Fig 6A-1, red box). A movie of the 3D reconstruction for HYPE density obtained from the tilt series is shown in Fig S6-Movie1, and a representative single slice from this movie is shown in Fig 6A-2. The thresholded density for HYPE (colorized in maroon) in this movie is presented in Fig S6-Movie2 and as a representative single slice in Fig 6A-3. These reconstructions confirm the periodic distribution of HYPE density throughout the lumenal face of the ER membrane.

**Figure 6.**
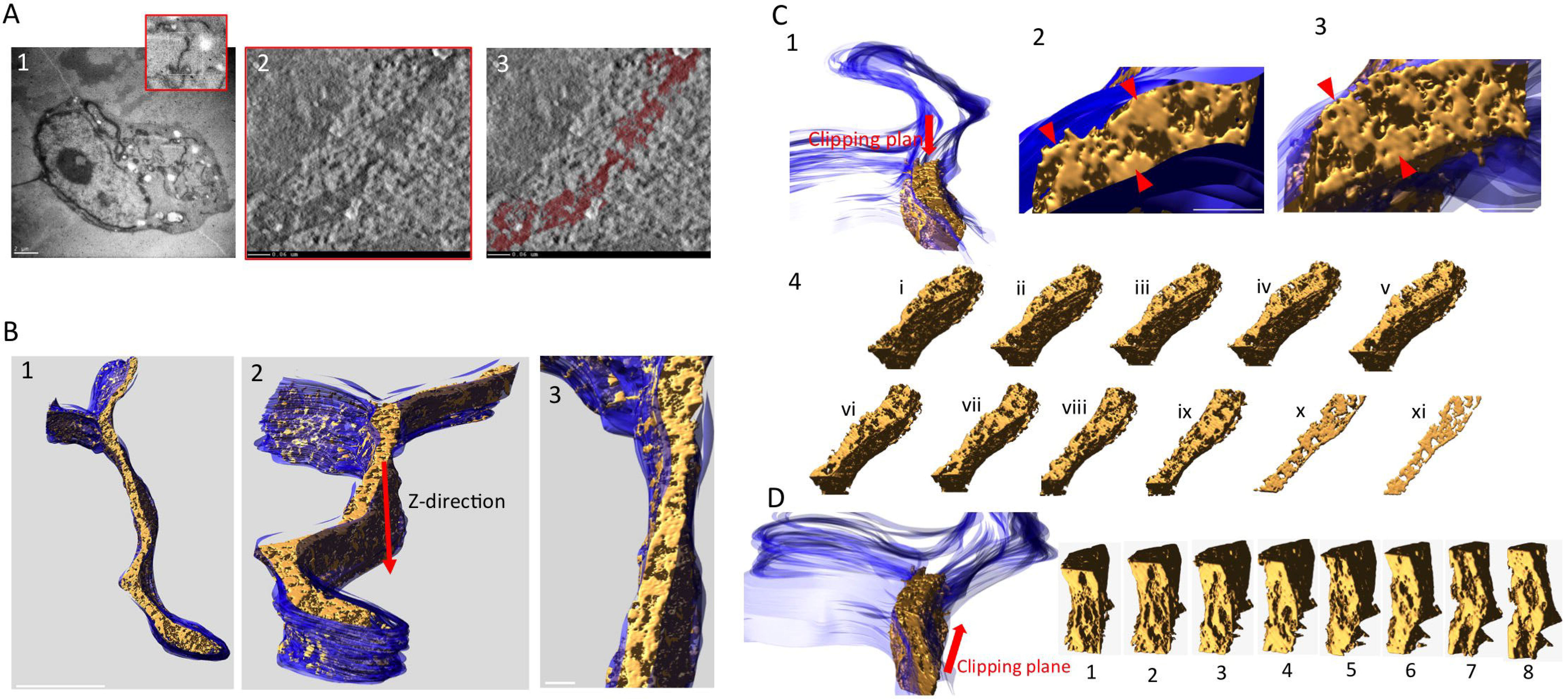
EM tomographic reconstruction of the ER exhibiting HYPE-APEX2 density. Tilt-series from HEK-293T cells expressing HYPE-APEX2 and processed by cryoAPEX were collected for 3D reconstruction of HYPE specific density. **A)** An image of the whole HYPE-APEX2 expressing cell showing an area containing ER tubules from where the tilt-series was collected (A1 and magnified red box). **A2)** Snapshot from the movie (see supplemental data, Figure S6 - movie 1) showing the reconstructed tomogram of HYPE density within the ER lumen. **A3)** Snapshot from the movie (see supplemental data, Figure S6 - movie 2) showing the reconstructed tomogram of the thresholded HYPE density (in maroon). **B)** An additional view of the 3D model of the ER membrane (blue) and the HYPE density within (gold) generated and visualized with the IMOD “Isosurface” tool (B1 through B3). **C)** Snapshots of a segment of the ER showing the HYPE-APEX2 density (gold) modeled within the lumen of the ER membrane (blue) visualized from the top of the indicated clipping plane (C1 through C3). Magnified images of different regions within this segment show HYPE’s periodic density pattern along the lumenal walls (C2, red arrow heads). Another view exposing the full face of the density (gold) using visualization tools that make the membrane transparent are also shown (C3, red arrow heads). This pattern of HYPE specific density is more apparent when the clipping plane is moved downward in the Z-direction progressively shaving through the depths of the different layers (C4, slices i through xi), and is most apparent when visualized in the thinnest slice (slice xi). **D)** A clipping plane that moves in a head to tail direction shows HYPE’s density pattern on the ER membrane from a front-on perspective (slices 1 through 8).

Next, 3D modeling by segmentation and visualization of the HYPE-APEX2 density showed that HYPE was confined within the ER membrane (Fig 6B). We further assessed this reconstructed segment of the ER from different visual perspectives (Fig 6B-1, whole ER segment; 6B-2, Z plane; and 6B-3, top down). Finally, we used the “clipping planes” tool to incrementally trim off the HYPE associated density from the top to the bottom (moving in the z-direction; Fig 6C and slices i through xi in Fig 6C-4) and in a head-to-tail direction (Fig 6D, slices 1 through 8). Both clipping planes substantiated our observation of the periodic foci for HYPE on the lumenal walls of the ER (Figs. 6C-2 and 6C-3, red arrows). Thinner, virtual slices of this region resembled the density pattern seen in 2D imaging of thin sections (compare Fig. 6C-4, slice xi with Fig 3C).

## Discussion

Fic-mediated adenylylation/AMPylation is an important, evolutionarily conserved mechanism of signal transduction. In humans, AMPylation mediated by HYPE regulates the unfolded protein response via reversible modification of BiP *(Ham et al., 2014; Preissler, et al., 2017; Sanyal et al., 2015)*. It is an open question as to whether HYPE functions beyond its role as a UPR regulator with additional physiological targets. Indeed, despite the fact that we and others have previously shown that HYPE is an ER membrane protein that faces the lumen, a number of candidate targets have recently emerged for further evaluation that reside outside the ER *(Broncel et al., 2016; Truttmann et al., 2016)*. For instance, the *C. elegans* homolog of HYPE (Fic-1) can be detected in the cytosol and is implicated in controlling the function of cytosolic chaperones *(Truttmann et al., 2017)*. HYPE’s *D. melanogaster* homolog (dFic) is implicated in blindness and shown to associate with cell surface neuro-glial junctions possibly by entering the secretory pathway *(Rahman et al., 2012)*. Thus, a clear subcellular localization for HYPE is needed to better understand its role in the context of these new protein targets - which led us to develop a technique to determine the subcellular localization of membrane proteins like HYPE at a high (low-nanoscale) resolution.

Despite tremendous advances in light microscopy, electron microscopy still remains the technique of choice to visualize cellular ultrastructure or determine protein localization at nanoscale resolution. Here, we describe the development of an EM method, called cryoAPEX, that successfully adapts the APEX2 tag for cryofixation while simultaneously retaining membrane preservation. Additionally, we show that data obtained by cryoAPEX for visualizing ER proteins, HYPE and ERM, can be used for EM tomographic reconstruction of membranes in 3D. Applying cryoAPEX to HYPE localization, we show that HYPE appears to reside solely in the ER lumen and in the contiguous nuclear envelope, in agreement with immunofluorescence data from us and others *(Sanyal et al., 2015; Truttmann et al., 2015)*.

CryoAPEX is designed specifically for localizing membrane-bound proteins. The methodology we present here enables sufficient resolution and membrane preservation such that even structures in close contact with the ER, such as the Golgi, mitochondria, or plasma membrane, are clearly distinguishable as HYPE-negative (Figs. 4-2, 4-3, and 4-4). These results tend to argue against the presence of a secreted or nuclear (other than the nuclear envelope) form of HYPE. However, any trace amounts of HYPE that may potentially correspond to processed or soluble isoforms that may localize to the cytosol would not be detected unless present as immobilized complexes. We also cannot rule out the possibility that trace amounts of HYPE could exist in subcellular compartments other than the ER, or that specific stimuli could result in a dramatic reorganization of HYPE to other locations. Still, if this were the case, we predict that by overexpressing HYPE relative to endogenous levels (as we do in our study), we would see HYPE more broadly distributed throughout the cell or perhaps even mislocalized. Our data show that even upon overexpression, HYPE is most robustly observed solely within different domains of the ER, at least under resting conditions. Experiments to address the possibility of HYPE’s relocalization in response to certain physiological stress signals are currently underway.

Intriguingly, our cryoAPEX analysis indicates that HYPE localizes as periodic membrane foci spanning the lumenal wall of the ER. The reasons are unknown, but this indicates that HYPE is tethered possibly via its hydrophobic N-terminus to repetitive structural elements along the ER, potentially as part of a larger signaling complex. The crystal structure of HYPE indicates that it exists as a dimer, interacting via its Fic domain *(Bunney et al., 2014)*. Additionally, the activity of bacterial Fic proteins, including HYPE, has been shown to undergo regulation by transitioning between monomeric, dimeric and oligomeric states *(Dedic et al., 2016; Stanger et al., 2016; Casey et al., 2017)*. Thus, it is possible that the HYPE specific periodic densities that we observe may represent HYPE oligomers. Further, we were able to reconstruct this HYPE specific density in 3D along the ER lumenal membrane using TEM tomography (Figure 6).

In our quest for the perfect EM based localization technology to visualize HYPE in the context of the ER and the whole cell, we were cognizant of the fact that to get a reliable picture of the presence or absence of HYPE in various subcellular compartments, we would require impeccable ultrastructural membrane preservation. Many organelles and transport vesicles within a cell are labile structures that are difficult to preserve in their native morphology. An organelle like the ER has multiple domains that make contacts with several other organelles as well as the plasma membrane. The functional relevance of these organellar contact points is of intense research and, in the case of the ER, they are known to be portals of lipid and calcium transport *(English & Voeltz, 2013; Rowland & Voeltz, 2012)*. Thus, preservation of these structures was especially important for ascertaining the distribution of HYPE.

Next, we considered the applicability of various traditional protein localization techniques such as immunoelectron microscopy (IEM), metal-tagging EM (METTEM) and peroxidase tagging. Unfortunately, each of these techniques suffers from a variety of limitations in addition to inadequate sample preservation. Specifically, current methods of detection are based on two common processes - 1) a chemical fixation step that precedes the actual detection assay and 2) a sample preparation step involving dehydration of fixed cells via alcohol at room temperature or on ice. This combination of chemical reagents leads to poor preservation of the membrane morphology due to lipid extraction and introduces artefacts. Therefore, in addition to membrane preservation, the method of choice needs to be compatible with heavy metal staining, so as to impart an adequate level of contrast between HYPE-associated membranes and other organellar membranes for contextual information about the ultrastructural environment within which HYPE resides. This ruled out METTEM tagging, as the technique suffers from its lack of compatibility with the use of heavy metal stains *(Risco et al., 2012)*.

Lastly, we considered the amenability of the method to 3D electron microscopic techniques. This is important as organelles or membrane structures such as the ER, Golgi, mitochondria, or the plasma membrane cover a vast 3-dimensional subcellular space and are in a constant state of morphological equilibrium with their surroundings. They undergo constant remodeling in their different domains in response to functional cues that can alter the localization of proteins that are associated with them *(Shibata et al., 2010; Voeltz et al., 2002)*. To detect such changes or, alternatively, the exclusive localization of a target protein in specific domains of these large organelles, 3D information at the site of protein localization can yield critical clues about protein function. Thus, we opted to develop a method that incorporated each of the above criteria to yield HYPE’s subcellular distribution in an optimally preserved and 3D EM compatible sample.

Cryofixation of live cells under high pressure (HPF) is a method that shows the best ultrastructural preservation and is now routinely used to prepare samples for EM tomography *(McDonald et al., 2006; O’Toole et al., 2018)*. It is not deemed compatible with most of the detection methods described above, however, as they require chemical fixation. Thus, to determine the subcellular distribution of HYPE, we developed cryoAPEX, a hybrid method that combines the power of APEX2 genetic tagging and HPF cryofixation. Chemically fixed cells expressing APEX2 tagged HYPE were first reacted with DAB to generate HYPE specific density, and then cryofixed and freeze substituted with acetone. As shown, cryoAPEX not only displays specificity of detection at high resolution for both lumen-facing (HYPE) and cytosol-facing (ERM) proteins, but also retains ultrastructural preservation that makes cryoAPEX amenable to TEM tomography. Further, cryoAPEX can be used to assess cells grown in monolayers making it widely applicable.

An important aspect of cryoAPEX is the robustness of the DAB byproduct that can withstand a long freeze substitution reaction in acetone. We also noted that cells once chemically fixed and labeled with DAB do not need to be cryofixed right away. In our hands, aldehyde fixed, DAB labeled cells that were cryofixed after 48 hours (storage at 4°C) exhibited no deterioration of cellular ultrastructure and staining when compared to those that were cryofixed immediately. This feature could prove to be of great advantage to laboratories that do not have immediate access to HPF and freeze substitution units. While this manuscript was being prepared, a technique with a similar goal of combining cryofixation with an APEX2 based detection method for carrying out CLEM (correlative light-EM) at the tissue (versus subcellular) level was reported. This method, called CryoChem, is procedurally different from our method. It requires immediate cryofixation of tissues and entails elaborate rehydration and dehydration steps following the initial freeze substitution in order to obtain better temporal resolution for CLEM studies *(Tsang et al., 2018)*. Also, CryoChem is yet to be tested on cell monolayers.

In conclusion, we have developed cryoAPEX as a method for obtaining localization of a single APEX tagged protein at a high resolution while maintaining excellent ultrastructural preservation and compatibility with EM tomography. We applied cryoAPEX to assess the subcellular localization of the human Fic protein, HYPE, and show that it is robustly detected very specifically on the lumenal face of the ER membrane and in cellular compartments that are contiguous with the ER lumen, where it displays periodic distribution resembling possible signaling complexes. Further, we do not detect HYPE in the mitochondrion, nucleus, plasma membrane or the Golgi/secretory network at the expression levels tested. Additionally, we show that cryoAPEX works equally well for cytosol-facing membrane proteins, like ERM, and accurately reflects ultrastructural morphological changes. Finally, we demonstrate that cryoAPEX can be applied to assessing protein localization using cell monolayers and executed in basic cell biology laboratories with relative ease.

## Materials and methods

### Plasmids

The HYPE-APEX2 plasmid was ordered through Genscript. Briefly, the HYPE-APEX2 fusion was first synthesized and then inserted into pcDNA3 vector between BamHI and XhoI sites. ERM-APEX2 plasmid was obtained from Addgene (plasmid no: 79055) https://www.addgene.org/79055/. The mito-V5-APEX2 was obtained from Addgene, (Plasmid#72480); CAAX-APEX2 was a kind gift from Alice Y. Ting and the ManII-APEX2 was ordered through Epoch biosciences. Briefly, a fusion construct comprising of the first 118 amino acids of the mouse isoform of α-mannosidase with the APEX2 gene in its C-terminus following a short intervening linker sequence was first synthesized and then cloned into pcDNA3.1. The complete sequence for MannII-APEX2 is provided (Supplemental data 1).

### Transfection and chemical fixation

HEK-293T cells were grown in 10 cm dishes and transfected with APEX2-tagged mammalian expression plasmids using Lipofectamine 3000 (Thermo Fisher). Cells were washed off the plate 12-15 hours post-transfection with Dulbecco’s PBS and then pelleted at 500x g. For those samples requiring chemical fixation, pellets were resuspended in 0.1% sodium cacodylate buffer containing 2% glutaraldehyde for 30 minutes, washed 3x for 5 minutes with 0.1% sodium cacodylate buffer and 1x with cacodylate buffer containing 1mg/ml 3,3’-Diaminobenzidine (DAB) (Sigma-Aldrich). Pellets were then incubated for 30 minutes in a freshly made solution of 1mg/ml DAB and 5.88mM hydrogen peroxide in cacodylate buffer, pelleted and washed 2x for 5 minutes in cacodylate buffer and once with DMEM. Finally, cell pellets were resuspended in DMEM containing 10% FBS and 15-20% BSA and pelleted. The supernatant was aspirated and excess media wicked off with a Kimwipe in order to remove as much liquid as possible. For BHK controls, cells were either cryofixed directly or prefixed with glutaraldehyde prior to cryofixation. An identical freeze substitution protocol was used for processing for both HEK and BHK cells. BHK cells were post stained with uranyl acetate and Sato’s lead prior to imaging.

### High pressure freezing and freeze substitution

Cell pellets (2-3 μl) were loaded onto copper membrane carriers (1mm x 0.5 mm; Ted Pella Inc.) and cryofixed using the EM PACT2 high pressure freezer (Leica). Cryofixed cells were then processed by freeze substitution using an AFS2 automated freeze substitution unit (Leica). An extended freeze substitution protocol was optimized for the preferential osmication of the peroxidase-DAB byproduct. Briefly, frozen pellets were incubated for 24 hours at -90oC in acetone containing 0.2% tannic acid and then washed 3x for 5 minutes with glass distilled acetone (EM Sciences). Pellets were resuspended in acetone containing 5% water and 1% osmium tetroxide, with or without 2% uranyl acetate (as applicable) for 72 hours at -80°C. Following this extended osmication cycle, pellets were warmed to 0°C, over 12-18 hours. Pellets were then washed 3x for 30 minutes with freshly opened glass distilled acetone. Resin exchange was carried out by infiltrating the sample with a gradually increasing concentration of Durcupan ACM resin (Sigma-Aldrich) as follows: 2%, 4% and 8% for 2 hours each and then 15%, 30%, 60%, 90%, 100% and 100% + component C for 4 hours each. Resin infiltrated samples in membrane carriers were then embedded in resin blocks and polymerized at 60°C for 36 hours. Post-hardening, planchets were extracted by dabbing liquid nitrogen on the membrane carriers and using a razor to resect them out of the hardened resin. After extraction of membrane carriers, a thin layer of Durcupan 100+C was added on top of the exposed samples and incubated in an oven for 24-36 hours to obtain the final hardened sample blocks for sectioning.

### Sample preparation via conventional room temperature method

HEK-293T cells were grown on collagen coated glass coverslips and transfected with HYPE-APEX2 or ERM-APEX2 mammalian expression plasmids for 24 hours as above. Cells were then washed with DPBS and chemically fixed with 2% glutaraldehyde in 0.1% sodium cacodylate buffer for 30 minutes. Fixed cells were washed with cacodylate buffer and finally with 1mg/ml of DAB in cacodylate buffer for 2 minutes. Following the wash, cells were incubated for 30 minutes in a freshly made solution of 1mg/ml of DAB and 5.88 mM of hydrogen peroxide in cacodylate buffer at room temperature. Cells were washed 3x for 2 minutes each with DPBS, incubated in an aqueous solution of 1% osmium tetroxide for 10-15 minutes and then washed with distilled water. Dehydration was carried out using increasing concentration of 200 proof ethanol (30%, 50%, 70%, 90%, 95%, 100%) followed by resin infiltration of the cells with gradually increasing concentrations of Durcupan resin in ethanol (30%, 60%, 90%, 100%, 100%+C). Coverslips were placed on beem-capsules filled with 100% + C cell-face down on the resin and incubated in an oven for 48 hours at 60°C. After polymerization, coverslips were extracted by dipping the coverslip-face of the blocks briefly in liquid nitrogen. Serial sections were then obtained by sectioning of the blocks *en face* and ribbons collected on formvar coated slot grids.

### Serial sectioning, lead staining and electron microcopy

Thin (90nm) serial sections were obtained using a UC7 ultramicrotome (Leica) and collected onto Formvar coated copper slot grids (EM sciences). Glass knives were freshly prepared from glass sticks during each sectioning exercise. Lead staining of the sections was carried out for 1 minute wherever applicable with freshly made Sato’s lead solution. Samples were screened on a 80 KV (Technai T-12) transmission electron microscope, and an average of 15-20 cells from multiple blocks were visualized for each sample.

### Measurement of HYPE-APEX2 density foci

The ImageJ software was used to measure the HYPE-APEX2 density foci within the ER lumen. Briefly, using the line selection tool the image scale was set up using the embedded scale bar as a yardstick. From the menu command, the “analyze/set scale” was used to set up the scale by entering the dimensions of the raw image (length of the embedded scale bar) in the “known distance” box. The unit of the scale bar was then set in the “units of the length box”. Readings were taken from multiple ER tubules and the range and average calculated.

### EM tomography, data reconstruction and segmentation

Thicker (250 nm) sections were used for collecting tomographic tilt-series. Sections were coated with gold fiducials measuring 20nm in diameter prior to collection. Tilt-series of a single 250nm section were collected with automation using the program SerialEM *(Mastronarde, 2003)* on a JEOL 3200 TEM operating at 300kV. The collected tilt-series were then aligned and tomogram generated by weighted back projection using the eTomo interface of IMOD *(Kremer et al., 1996)*. The reconstructed tomogram was visualized in IMOD. The ER membrane was first hand segmented and then used as a mask for thresholding of the density within the ER lumen.

## Supplemental Material

Supplemental materials includes Figure S1, Figure S2, Figure S3, Figure S4, Figure S5, and two movies accompanying Figure 6, snapshots of which are represented in Figure S6. The two movies are included as separate movie files labeled Sengupta & Mattoo - CryoAPEX JCB - Fig S6 Movie 1 and Sengupta & Mattoo - CryoAPEX JCB - Fig S6 Movie 2.

## Acknowledgements

We thank Dr. Jason Lanman (Purdue University) for access to HPF, FS, and microtome units. We are also grateful to Drs. Alice Ting and Jeffrey Martell (Stanford University) for advice and providing the CAAX-APEX2 (*Lam et al., 2015*) construct and to Dr. Ben Giepmans (University of Groningen) for providing their Golgi-FLIPPER (*Kuipers et al., 2015*) construct and sequence from which we cloned our mannII-APEX2 construct. We are indebted to Ms. Elaine Mihelc for help with IMOD segmentation and technical discussions, and to Dr. Robert Stahelin for constructive and thoughtful discussions. We are also grateful to members of the Mattoo lab for feedback. Thin section screening was carried out at the Purdue Life Sciences EM Core Facility. Tomogram collection was conducted at the Electron Microscopy Center at Indiana University – Bloomington under guidance from Dr. David Morgan.

## Author contributions

SM conceived and supervised the study. RS conducted the experiments. RS & SM designed and developed the study, and analyzed the data. SM, MJP, and RS wrote the manuscript.

